# Simple bacterial growth model for the formation of spontaneous and triggered dormant subpopulations

**DOI:** 10.1101/2023.08.18.553832

**Authors:** Mikkel Skjoldan Svenningsen, Namiko Mitarai

## Abstract

Bacterial persistence is a phenomenon where a subpopulation of cells can survive antibiotic treatment, and it is often linked to extremely slow growth or a dormant state. However, the mechanisms and factors that govern dormancy are not well understood. We propose a simplified growth model that treats the main cellular components as discrete variables and allocates resources among them according to different strategies. The model can reproduce some of the observed features of bacterial persistence, such as wide distribution in division times, long division times after a nutrient down-shift, and the existence of different dormant phenotypes. We also show how the growth structure, i.e., whether cells grow in a lineage or a branch, affects the dormant cells’ occurrence and distribution due to the growth states’ mother-daughter correlation. Our model provides a framework to explore the complex interactions between cellular processes and environmental conditions that lead to bacterial persistence.

## I. INTRODUCTION

When a lethal amount of antibiotic is applied to a bacterial population, the majority is killed exponentially over time [1, 2]. However, there is almost always a minority of cells in the antibiotic-sensitive population that dies at slower rates [3–6]. These cells are generally referred to as bacterial persisters [7]. The persisters seem to play a key role in chronic infections by surviving antibiotic treatments and later giving rise to new infections [8]. In addition, persistent cells were linked to the rise of resistant cells [9]. Despite the massive impact on human health and the discovery of this cellular state more than 70 years ago [3], the molecular mechanism of persister formation is not yet fully understood. Bacterial persistence appears to be a complex phenomenon that involves various timescales [6], and thus, it is likely that the bacterial persister population is comprised of multiple different subpopulations.

Bacterial persistence is often associated with slow or nongrowing cells. In fact, since the discovery of persister cells, they have been linked to dormancy [4, 10–12]. A naïve interpretation of the link is that most types of antibiotics target cellular growth processes, hence they are not lethal for dormant cells. With this interpretation, the killing curve reflects the distribution of dormancy duration [13], i.e., if a bacterium resumes growing under antibiotic application, it will be killed. Given the association between dormancy and persistence, understanding the cell-to-cell distribution of doubling times, including the formation of extremely slow-growing cells in a faster-growing population, could help understand bacterial persistence.

Doubling time, or interdivision time, distributions have been shown to have functional forms close to a Gaussian or a gamma distribution [14–16]. Those distributions have similar shapes close to the mean but differ in the tails. However, the tails of the interdivision time distribution are difficult to determine experimentally due to the necessity of large statistics. Yet, it should be noted that the tails from both a Gaussian and a gamma distribution predict a decay equal to or faster than an exponential. In contrast, the killing curves of the persister population suggest that the tail of the doubling time distribution decays slower than the single exponential extrapolated from the majority of the population. This indicates that persistent cells may represent subgroups in different physiological states than the main population.

The molecular mechanisms of bacterial persistence are not understood yet, but it is very likely that there are multiple different mechanisms. Some of the mechanisms are linked to specific sets of genes, such as stochastic activation of toxin-antitoxin system [17] and unequal expression of efflux pumps [18]; some of them may not even be linked to slow growth or dormancy [19, 20].

However, other proposed persistence mechanisms could potentially be understood from simple growth principles. Radzikowski et al. hypothesized that persisters enter a state of dormancy due to collapses in the central metabolism, leading to a state of low intracellular nutrient levels [21]. The assumption is that a positive feedback loop in the metabolic network can lead to some cells entering a dormant state and thus surviving antibiotics. Another study has demonstrated how dormancy could arise as physiological states from the kinetics of the bacterial metabolic network [22]. In addition, it was previously demonstrated that a clonal population of bacteria can exhibit switch behaviour in nutrient uptake [23, 24]. Several studies link low ATP levels to bacterial persistence [12], which could support the idea of collapses in the central metabolism and growth deficiencies in general.

Such a view on persistence may also enable us to understand the spontaneous persistence in the steady-state growth and the triggered persistence when cells are exposed to external stress such as starvation [7] in a coherent manner. One possible scenario is that the spontaneous persisters are the cells that experience the collapse of the growth process by inherent ffluctuations in the biochemical processes, and the transient stress pushes the growth process closer to collapse, triggering more cells to enter dormancy that cannot be immediately recovered when the stressor is gone.

To understand the possible origins of heavy tails in survival time distributions, we propose a growth model based on general growth principles of bacterial physiology in a manner inspired by recent approaches to coarsegrained growth models [25, 26]. A bacterial genome is remarkably optimized for growth across varying conditions, where the cell must allocate resources between protein synthesis, cell division, energy production, amino acids synthesis, nucleotides, etc. The allocation problem is solved in a noisy environment, with both intrinsic and extrinsic sources of noise [27], leading to heterogeneity in a clonal population. We construct a simple growth model that phenomenological takes into account the resource allocation and the noise. We aim to demonstrate how these growth principles could lead to extremely long doubling times in both steady-state growth and under external temporal stress. We then discuss the relation between our findings and the experimental results, especially our recent experiment on spontaneous and triggered persistence in a controlled environment [6], to provide a base for future theoretical considerations on bacterial growth heterogeneity and persister formation.

## II. THE MODEL

### A. Stochastic growth model

We built a mathematical model for cellular growth and division based on simple principles illustrated in Fig. 1. The cell imports nutrients and uses them for three different processes: building translation machinery, facilitating the cell division and importing more nutrients. These sectors are represented by the variables *n* (intracellular nutrient level), *r* (translation), *s* (division), and *m* (nutrient uptake). The model is highly coarse-grained, and each variable represents a sector of content, rather than specific proteins, inspired by previous models of bacterial physiology [28, 29].

**FIG. 1.**
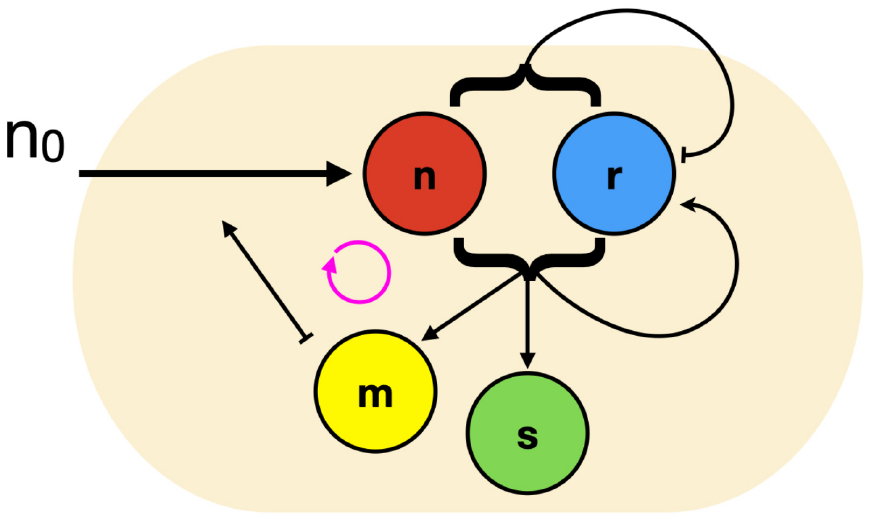
Illustration of mathematical model showing the main features of the regulatory network. The *n*_0_ quantifies the extracellular nutrient concentration. The uptake of nutrients is dependent on *m*. All variables, except for *n*, are dependent on the combination of *n* and *r* for production. The production of *r* is regulated according to both the level of *r* and *n*.

The variable *r* resembles the number of ribosomes per cell but generally refers to the translational machinery, which was demonstrated to be concurrently regulated [29, 30]. The variable *s* refers to the cell division machinery. The bacterial division is complex and results from the cooperation of several proteins. In fact, the exact mechanism of cell division in *Escherichia coli* is not known yet [31]. Inspired by a model of the adder principle [15], the variable *s* represents the general division sector in our model, and once a sufficient amount is allocated, the cell divides. The division limit is set by the parameter *s*_0_, which is set to be an even number; just after a division, each daughter cell gets *s*_0_*/*2 units of *s*, and the next division happens after *s*_0_*/*2 units more of *s* are produced. The variable *m* quantifies the cell’s ability to import and metabolize exogenous nutrients. This coarse-grained variable includes the effects of, e.g., in the case of *E. coli*, such as the nutrient uptake by the passive diffusion through the outer membrane channels and the active uptake by the phosphotransferase system [32], as well as the biosynthesis of necessary amino acids from imported nutrients. The intracellular nutrient level *n* is also a coarse-grained variable representing the materials needed to build all the necessary machinery for cellular growth. The variables *r, s* and *m* are all exclusively decaying by dilution when the cell divides. The variable *n* decays both by consumption and dilution.

The model is implemented as a stochastic process using the Gillespie algorithm [33] with the rates shown in Table I combined with a jump process for cell division: Every time the variable *s* reaches the threshold *s*_0_, the cell divides and two new cells are produced with half the cellular content in each (if the content is an odd number, the content is randomly divided such that one cell gets one more unit than the other cell). All variables are discrete and refer to the integer number of content per cell. However, due to the coarse-grained nature of the model, they do not represent the number of molecules per cell. Rather, they are the units that are collectively needed to perform the task, and also their number determines the noise level in our simulation.

**TABLE I.** The mathematical model implemented as a Gillespie simulation. For each event, there is a given rate. The functional forms are 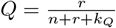 and 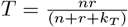.

| Event | Variable update | Rates |
| --- | --- | --- |
| 1. nutrient uptake | $n \rightarrow n + 1$ | $\xi + n_0 m$ |
| 2. $r$ production | $n \rightarrow n - 1, r \rightarrow r + 1$ | $\epsilon n + \omega(1 - Q) \cdot T$ |
| 3. $s$ production | $n \rightarrow n - 1, s \rightarrow s + 1$ | $(1 - \phi)QT$ |
| 4. $m$ production | $n \rightarrow n - 1, m \rightarrow m + 1$ | $\phi QT$ |

The content of each cell in the growing population is simulated explicitly to analyze the cell-to-cell variability of growth rates and their possible mechanistic origins. This is in contrast to the growth models where cellular content is either modelled as the time-evolution of the probability density for cellular content [26] or simply the population mean [25]. It is worth mentioning that a stochastic version of a model that considered cellular content at a single cell-level was analyzed in [34] to obtain the growth rate and size distribution in an exponentially growing population, focusing on the behaviour around the major peak of the distribution. Here, we focus on a larger noise level that produces dormant subpopulations at a detectable level.

The functional dependence of the rates on the variables is kept as simple as possible, as seen in Table I. The nutrient uptake depends linearly on the variable *m*, plus a very small constant infflux *ξ* of nutrients. Here, *ξ* represents the infflux of building blocks without the help of explicit metabolic proteins. The translational capacity is given by the function 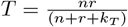, corresponding to the cell’s ability to make either *r, s*, or *m* per unit of time. The numerator of *T* is the product of *n* and *r* because the translational capacity depends on both the translational machinery and energy/nutrients.

Because of the coarse-grained nature of the model, the correspondence between the simulation units and the real units can be inferred only through the resulting behaviour. As we see in Fig. 2A the fastest growth rate with the saturating amount of nutrients is 1 for the default parameter sets. Then, for example, for *E. coli*, the maximum doubling time is about 20 min, making the simulation time unit to be about 0.5 hours. The conversion is harder for the concentration unit because one unit of different variables is likely to correspond to different protein mass.

**FIG. 2.**
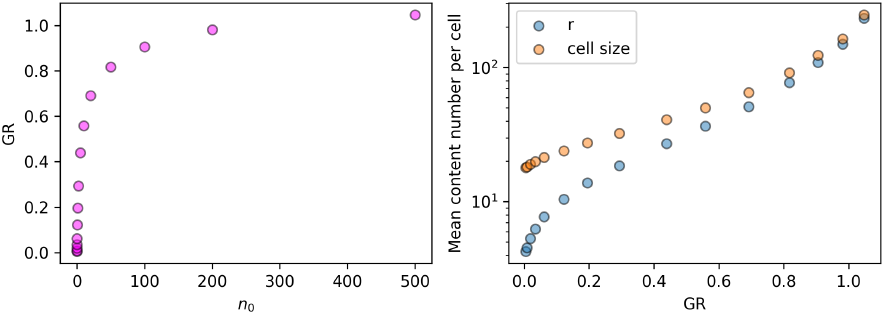
Growth-law tendencies reproduced by the growth model. (A) The growth rate increases with the extracellular nutrient concentration to saturate at the maximum growth rate. (B) Both the cell size and the ribosome level per cell (*r*) increase with the growth rate.

Resource allocation is the core feature of the model. There are different ways to model resource allocation; for example, Erickson *et al*. [35] uses ffluxes of nutrients/metabolic molecules and their balances to determine the resource allocation, instead of explicitly modelling the nutrient concentration in a cell. However, inspired by refs. [25, 26], we assume that the allocation is explicitly dependent on the amount of nutrient available in the cell, governed by a dimensionless function 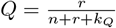 *Q* determines the allocation between ribosome production and the other sectors. The low/high value of *Q* represents the state where a large/small amount of resources is currently allocated for the translational machinery compared to the nutrient availability. At high nutrient availability, the majority of resources should be directed towards ribosome production to keep a high translation rate. At low nutrient levels, the resources should be directed more toward other sectors which include nutrient transporters and the division-associated sector. This consideration led to the model where the fraction (1 *− Q*) of the translational capacity *T* is devoted to the production of *r* and the fraction *Q* is devoted to the production of *s* and *m*.

Further allocation between the metabolic sector *m* and the division associated sector *s* is quantified by a constant parameter *ϕ*. The production rate of the *r* sector can be different from that of the *s* and *m* sectors, which is parameterized by *ω. ω* characterises the difference between one unit of ribosomal sector and one unit of other sectors: If one unit of ribosomal sector protein is twice as large in terms of molecular weight, it will take on average twice as long to make one unit compared to making one unit of the other. Since our model is very coarse-grained, we do not have a concrete number for *ω*. Therefore, we test different values of *ω* in the following simulation. Finally, to prevent complete growth stalling when *r* = 0, we allow *r* to be produced with a rate *ϵn*, where *ϵ* is a small parameter. The proportionality to *n* is because the production of *r* consumes nutrient *n*.

The total set of parameters in this model is *n*_0_, *k*_*T*_, *k*_*Q*_, *ϕ, ω, ϵ, ξ* and *s*_0_. The parameter *n*_0_ quantifies the extracellular nutrient concentration and represents the uptake per unit time per unit transporter. The parameter *k*_*T*_ represents a scale of saturation for translation. For all simulations, this value is set equal to *k*_*T*_ = 10. The parameter *k*_*Q*_ is there to avoid the collapse of the function *Q* when *r* = *n* = 0 and chosen to be 10^*−*9^.The parameter *ϕ* represents an allocation between energy and division, whereas *ω* determines the maximal production rate of ribosomes. We set *ϕ* = 0.5 and *ω* = 1.0 unless otherwise mentioned. The parameter *s*_0_ corresponds to the ‘Division control parameter’ in [25], but works here at the single-cell level. A cell divides when the variable *s* reaches the threshold *s*_0_. At a division event, two new cells are produced, and the content is distributed equally to each cell. If the content of the mother cell is an uneven number, one cell gets an extra unit of this value by a random event.

It is important to note that the noise level is regulated by the parameter *s*_0_ in the current model, since *s*_0_ affects the mean level of all the cellular components. The lower the value of *s*_0_, the lower the value of all components. It was previously demonstrated that for cell-to-cell protein distributions, the coefficient of variation decreases as the mean protein number increases [36]. In reality, noise crucial to the growth dynamics is affected by the ffluctuations of various limiting factors and the amplification of small noise by nonlinear dynamics in the complex cell growth process. Instead of modelling complex noise sources, we choose the value of *s*_0_ so that the noise level is high enough to observe dormancy within the available computational time, and we focus on the qualitative features of the dormant cells’ appearance. Hence, the value is fixed at *s*_0_ = 10 for the main text. Changing this value is also explored further in the supplementary Fig. S1.

The model inherently contains the possibility of slowly-growing states. If *ξ* = *ϵ* = 0, *m* = *n* = 0 is an absorbing state where no further reaction is possible. *r* = 0 will also be a no-growth state as the cell is incapable of making *m, r*, or *s* any more, though nutrients will keep accumulating if *m >* 0. Hence, *m* = *r* = 0 is another complete absorbing state where no growth or accumulation of nutrients is possible. The nonzero values of *ξ* and *ϵ* make these states transient dormant states instead of inescapable non-growing states.

In reality, it is likely that the exit process is not a simple constant rate process but complex dynamics that give non-trivial exit time distributions [22, 37]. However, because how the cell exits dormant processes is fairly understudied compared to the exponential growth process, we chose to model it in the simplest possible way so that it is easy to see the effect of these parameters. Therefore, the focus of the current model is to qualitatively analyze the possible entry to the dormant state based on the simplified growth dynamics, and not the recovery from the dormant state.

The two exit rates, *ξ* and *ϵn*, affect the statistics of the dormant subpopulations but do not affect the qualitative results as long as the values of *ξ* and *ϵ* are small enough. In the current simulation, their values are fixed at *ξ* = 10^*−*3^ and *ϵ* = 10^*−*6^. Dormant states are expected as long as the probability for entering either *r* = 0 or *m* = *n* = 0 is practically nonzero within the scope of the simulations. In the following, we refer to three different dormant states: the *m-dormant* (*m* = *n* = 0), the *r-dormant* (*r* = 0), and the *mr-dormant* (*m* = *r* = 0).

### B. Population growth

The model presented here describes the time evolution of cellular content in a single cell and the cell division process. We need to choose a sampling method to generate a population of growing cells from the model. A simple choice is to follow a single lineage of cells by keeping only one of the two cells at cell division [25, 26, 38], and it corresponds to the experimental setup where a lineage is followed in a mother machine [38, 39]. Another choice is generating a branching process of dividing cells and following cells in all the branches in parallel, corresponding to bacterial growth in a flask. Here, we call the former lineage growth and the latter branch growth.

These two growth structures are known to result in different distributions of doubling times [14, 16, 38, 40]. The difference between a lineage growth and a branching process becomes pronounced when the formation of a dormant cell is positively correlated with its ancestor cell slowly dividing: In a lineage growth, a slowly growing cell will eventually divide to produce a dormant progeny, but in a batch culture, the growing population take over the statistics exponentially fast before the division of a slowly growing cell. For interested readers, the comparison of different sampling methods for the population with growth heterogeneity is thoroughly studied in ref. [38].

In the following, most simulations were done with lineage growth unless otherwise noted. Data used for analyses were collected after 1000 cell divisions from the initial cell, and up to 10^6^ to 10^7^ cell divisions, and multiple simulations were done for each condition. In lineage growth, the growth rate was determined from the mean interdivision time. As will be shown in the results, our model produces a dormancy state, which is dependent on the status of the ancestor cell. Therefore, we also analyse the branching process to demonstrate the difference between the two statistics. When we simulate a branching process, first we perform linage growth for 10^4^ divisions, and then the branching process is simulated for a fixed period of time (20 times the mean lineage inter-division time). The last state was then used for analyses. The growth rate was determined by repeatedly growing a population of cells from one cell to *N* cells, within a fixed time interval Δ*T* . The growth rate from one simulation was determined as log(*N*)*/*Δ*T* and the mean of these values is then found.

The code used are available at https://github.com/mssven/Bacterial-growth-model.

## III. RESULTS

### A. The model mean behaviour follows established growth laws

We aim to capture the basic trends of bacterial growth physiology in the current simple model. We first compare the population mean values with established growth laws to check this.

Figure 2A shows that the growth rate increases and saturates as a function of the external nutrient concentration *n*_0_. This is consistent with the Monod growth law, i.e., the growth rate increases with the exogenous sugar concentration for *E. coli* with a hyperbolic relationship [41].

Figure 2B depicts the growth rate dependence of the mean cellular value of *r* (blue circles) and the sum of protein-like sectors *r* + *s* + *m* (orange circles), where the latter can be interpreted as cell size in the current simple model. The growth rate is controlled by changing *n*_0_ (Fig. 2A). We see that both quantities are increasing with the mean growth rate. On the fast growth rate (faster than 0.7) the trend can be approximated as one exponential, and in the middle growth rate (between 0.3 and 0.7), it appears to be exponential dependence with another scale. Experimentally, it has been shown by Schaecter *et al*. [42] that as cells grow faster by change of the carbon source, their mean ribosome content and the cell size increases exponentially with the growth rate in the range of growth rate 0.4 to 2/hour. Our model follows this trend in a limited range of the external nutrient concentration *n*_0_. It is worth noting that our model does not consider DNA replication explicitly, while Schaechter–Maaløe–Kjeldgaard growth law is normally associated with the DNA replication and cell growth coupling [43].

Another well-known growth law is that the fraction of ribosomal protein mass in the total protein mass increases linearly with the growth rate [28, 42]. It is worth mentioning that the previously proposed simple growth models required somewhat complex feedback control to reproduce the ribosomal growth law [25, 44]. Because our simple model ignores a large part of the essential proteome, e.g., those that contribute to the cell wall production, the ribosome fraction increases in the model but does not show linear dependence. For interested readers, *r/*(*r* + *s* + *m*) is plotted as a function of the growth rate in a supplementary Fig S2.

### B. Formation of dormant subpopulation

Cells with extremely long doubling times form occasionally during growth along a lineage. This is shown in Fig 3 where the doubling time distributions for various values of *ϕ* is presented. The blue distributions show the doubling time of cells that did not enter any dormant states. The yellow, green, and red distributions show the m-dormant state (*m* = 0, *r* ≠ = 0), the r-dormant state (*r* = 0, *m* ≠ 0), and the mr-dormant state (*m* = *r* ≠ 0), respectively. Each dormant states have a significantly longer mean doubling time than the non-dormant case (blue distribution). We see that the dormant state is more frequent for a lower value of *ϕ* (Fig 3AB): For values of *ϕ* above approximately 0.6 (Fig 3C) we did not observe any dormant state within the simulated duration. This depicts the importance of *m*-sector proteins for dormancy since higher *ϕ* means the allocation of more resources to the *m*-sector proteins.

**FIG. 3.**
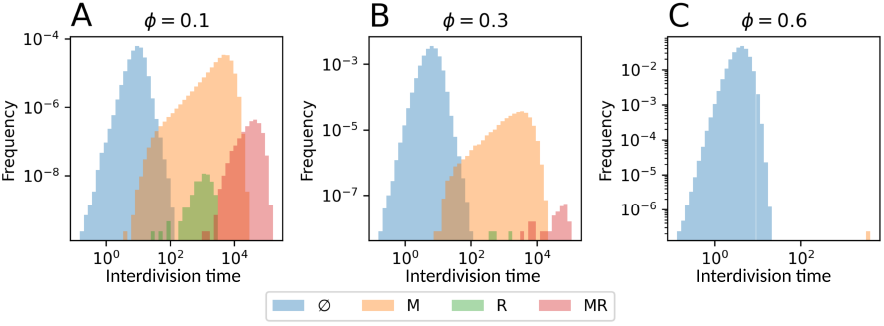
interdivision time distributions for three values of *ϕ*. Showing the fast-growing population (*∅*) and three dormancy types (*M*,*R*, and *MR*). The occurrence of the dormant types strongly depends on the parameter *ϕ*. The value of *n*_0_ was fixed to 1.0.

To further understand the role of resource allocation in the formation of dormant cells, we plot the mean value of *m, r*, or *n* for various values of *ϕ* and *ω* in Fig. 4A B, or C, respectively. The parameter *ϕ* determines the resource allocation between *m* (nutrient availability) and *s* (cell division), while the parameter *ω* controls the efficiency of nutrients converted to *r* proteins. The effect of these parameters on the *m*-, *r*-, and *mr*-dormant states as well as the mean growth rate are also shown in Fig. 4D E, R, and G, respectively. It is worth noting that the frequencies of dormant states do not simply anticorrelate with the mean growth rate (Fig. 4G). This is demonstrated further in the supplementary Fig. S3, showing the dormancy dependence on the population level growth rate. Increasing *ϕ* increases the mean value of *m* (Fig. 4A), and hence lowers the chance to enter the m-dormant states (Fig. 4D) due to a smaller probability of reaching *m* = 0. While the mean *m* level is independent of *ω*, we see a weak dependence of the m-dormant probability on *ω*. This is because low *ω* results in less *r*-proteins (Fig. 4B) hence slower consumption of the nutrients. This enables a cell to store energy (higher *n*) for low values of *ω* (Fig. 4C), reducing the probability to reach *n* = 0 which is also required for *m*-dormant state.

**FIG. 4.**
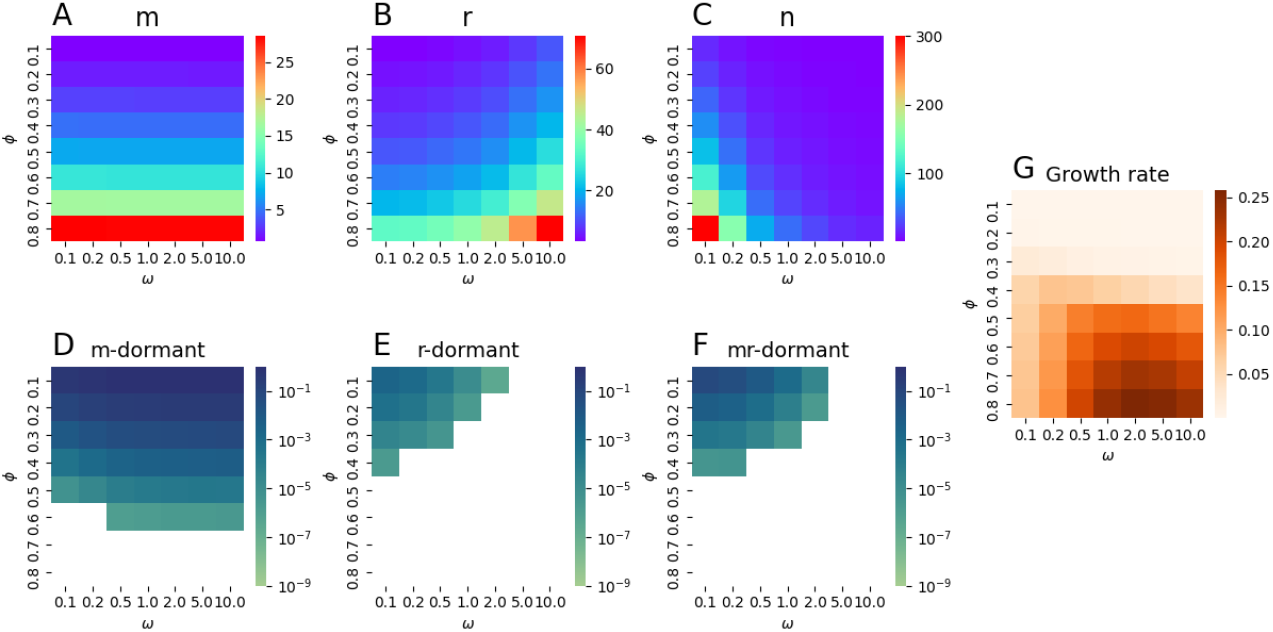
The dependence on resource allocation parameters *ϕ* and Ω. (A) mean value of *m*. (B) mean value of *r*. (C) mean value of *n*. (D) m-dormant state frequency. (E) r-dormant frequency. (F) mr-dormant frequency. (G) mean growth rate. *n*_0_ = 1 was used.

The mean value of *r* increases with *ϕ* and *ω* (Fig. 4B). Since the probability to reach *r* = 0 anti-correlate with the mean value of *r*, we naturally observe the opposite trend in the frequency of *r*- and *mr*-dormant states (Fig. 4EF). We observe that *mr*-dormant states are in general rarer than the *m*-dormant state and occur in a more limited part of the parameter space (Fig. 4E and F, respectively). In the next subsection, we show that the *r*- and *mr*-dormant states are preceded by *m*-dormant state, which explains the reason why the parameter region with higher levels of *r* and *mr*-dormancy are a subset of the parameter region with frequent *m*-dormancy.

### C. The interdependency of dormant states leads to a higher frequency of dormancy during lineage

**growth compared to branch growth**

The statistics presented so far were based on lineage growth. When we compared the frequency of dormant cells in lineage growth with the frequency in branch growth, we found a significant reduction in the branch growth (Fig. 5A). Noticeably, the *r*- and *mr*-dormant cells do not form during branch growth for this specific set of parameters within the simulation (the bars with hatch show the detection limit due to the sample size). The explanation for this difference is that there are mother-daughter correlations in the formation of dormant cells spanning several generations. Once a cell has entered a dormant state, the probability for the offspring to be dormant is strongly increased (see supplementary Fig. S1). In other words, if we follow a lineage, once a dormant state is observed, several generations of dormant states are likely to be observed. In contrast, in a branch growth, non-dormant cells take over the population before a dormant cell produces another cell that enters the dormancy.

**FIG. 5.**
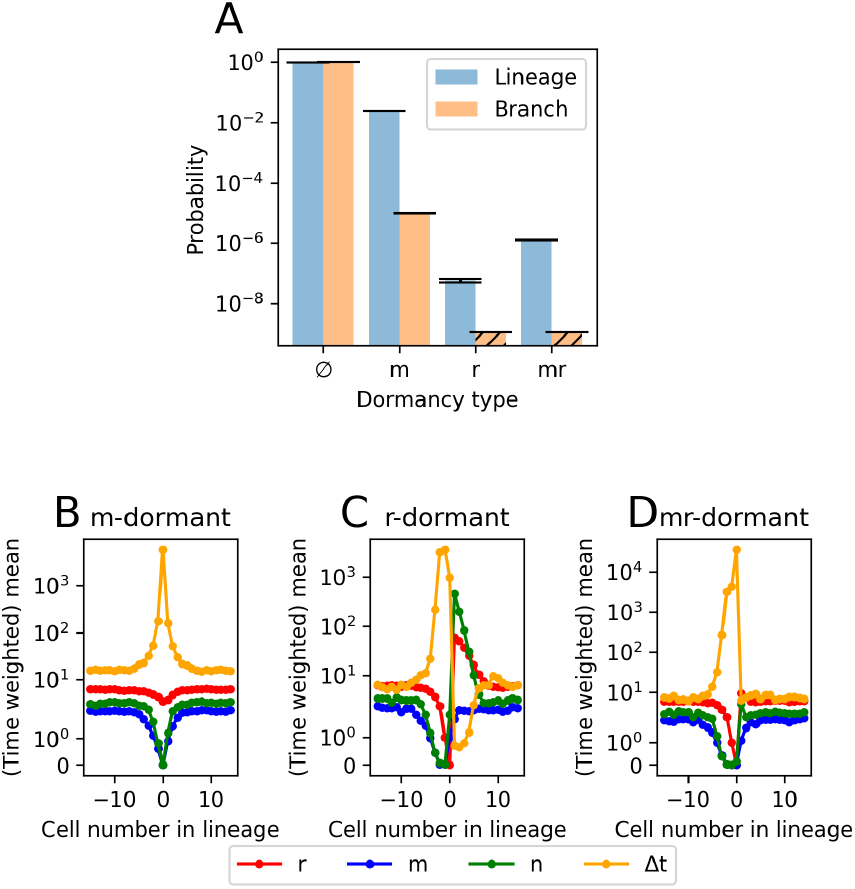
(A) Dormancy probability in the lineage growth (blue) and the branch growth (orange). The orange bars with hatch show the detection limit due to the sample size. (B) The time-averaged population mean of cellular content before, during, and after a dormant state along a linage. Each number corresponds to the cell number before and after entry into dormancy. Illustrating the necessity of *m*-dormancy in ancestry cells before entry into *mr*-dormancy. The yellow line is the interdivision time Δ*t*. For this simulation, *ϕ* = 0.3 were used.

To understand the correlation better, the mean content of cells leading up to and following a dormant state is depicted in Fig. 5BCD averaged over an ensemble of lineages. The cell assigned the number 0 is either *m*-, *r*-, or *mr*-dormant, respectively. The *m*-dormant state formation is least dependent on previous states, as seen in Fig. 5B in contrast to the other dormant states which are preceded by the *m*-dormant state (*m* = *n* = 0).

The formation of an *mr*-dormant state requires a few generations to form (Fig. 5D). We observe that the cell keeps dividing though at a slower rate when the nutrient starts limiting, indicating that the cell will keep producing *s* instead of *m*. As a consequence, *r* is kept diluted to eventually hit *r* = 0. Thus the probability for this to happen is an increasing function of 1*−ϕ*, since 1*−ϕ* is the rate to produce *s*. This is consistent with the observation in Fig. 4F.

Fig. 5C shows that the *m*-dormant state precedes the *r*-dormant state, but it enters the *r*-dormant state if the cell manages to produce *m* while *r* is diluted to be zero by cell division. Since *m* is non-zero, nutrient *n* can accumulate while the cell awaits the production of *r*. Furthermore, the resource will be fully allocated to *r*-production due to the functional form of *Q*. Hence, the cell just after the exit from r-dormancy can overshoot in the level of *r* and *n*.

### D. Nutritional downshift leads to a higher frequency of dormant cells in a manner dependent on the growth structure and allocation strategy

Next, motivated by the recent experiment on spontaneous and triggered persistence [6], we analyze the effect of a sudden nutritional downshift to the dormancy in our growth model. We model the transient nutritional downshift by setting *n*_0_ = 0 for a certain time interval and then returning the parameter to its original value. Here, the downshift lasts 10 units of time, which is well below the interdivision times of dormanct cells and at the same time well above fast-growing cells’ interdivision time. The aim is to investigate how a downshift affects the statistics of dormant cell formation and to determine how this effect is dependent on the model parameters.

We observed that the downshift strongly enhances the *m*-dormant fraction, but the effect is dependent on the parameter values. This is summarized in Fig. 6 where the fraction of *m*-dormant cells observed just after a downshift end (*P*_*ds*_) relative to the fraction of *m*-dormant cells without a downshift *(Pcontrol)*, 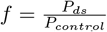, is plotted in the vertical axis. The downshift effect is very dependent on the mean level of *n* in the cells before the downshift, as seen in Fig. 6A. The downshift naturally leads to a lower level of *n*, and consequently shuts down the cell’s metabolism by reaching a *n* = 0 in some cells. The effect of the downshift is dependent on how the cell allocates its surplus of *n* before not being able to synthesise new content. As seen in Fig. 6A there is a threshold in mean *n* for the downshift effect, and the cell does not enter dormancy if the mean *n* is too low before starvation. This is because, to enter a *m*-dormant state, the cell should be able to keep dividing to dilute *m* to reach zero. The observed threshold value of *n* (about 15 units) is consistent with the observation that it requires a few cell divisions to dilute *m* enough, and it requires at least 5 units of *n* to divide once since *s*_0_ is set to 10.

**FIG. 6.**
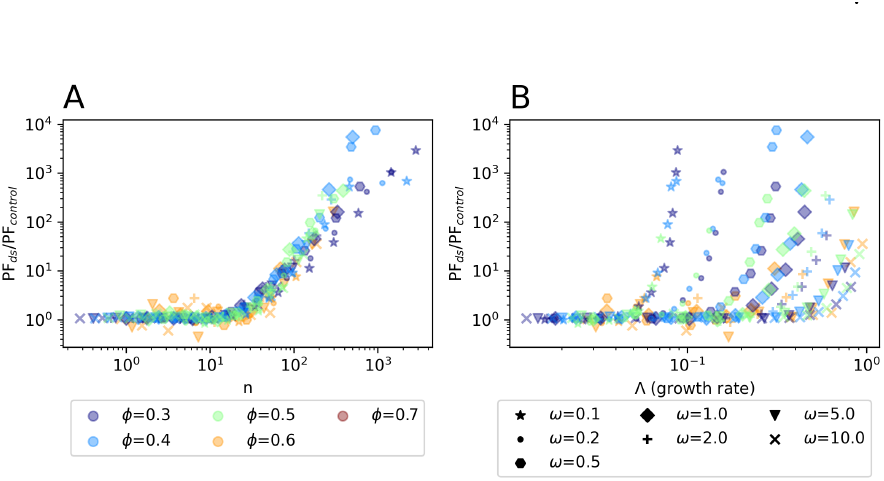
The effect of a transient downshift on m-dormancy formation. The fraction, given by *f* = *P*_*ds*_*/P*_*control*_, of dormant cells with and without a downshift is plotted as a function (A) the mean nutrient level in the cells or (B) the growth rate before downshift. The colour of the symbols represents the value of the *ϕ*, and the shape of the symbols representsthe value of *ω*. Different values of *n*_0_ were simulated for each parameter set to obtain different mean *n* values and growth rates.

We have also observed that the downshift effect correlates with the growth rate before downshift as shown in Fig. 6B in an *ω*-dependent manner. This can be understood through the mean *n* dependence presented in Fig. 6A: Because *ω* determines the efficiency of conversion from *n* to *r*, the lower values of *ω* result in less *r* and hence more storage of *n* in a cell (see also Fig. 4BC) for a given growth rate. Therefore, the cells with lower *ω* manage to divide more often during the downshift, resulting in a higher frequency of the dormant cells by reaching *m* = 0.

In nature, the bacteria are expected to experience cycles of nutrient-rich and starvation conditions repeatedly [45, 46]. We, therefore, also tested how the fraction of dormant cells changes when the nutrient availability *n*_0_ between a positive constant value and 0. We limited both the starvation and feast period to 10-time units (supplementary Fig. S4). Similar to the starvation pulse case of Fig. 6A a large difference with and without starvation period was observed only for relatively rich feast condition (*n*_0_ *≥* 10), where the fraction of *m*-dormants cell increased significantly.

## IV. DISCUSSION

The simplified growth model presented here attempts to explain a few observed features of bacterial persistence from simple principles to possibly understand some of the many factors at play rather than fully understanding the highly complex phenomena of interdivision time distribution and bacterial persistence. The proposed simple allocation model managed to mimic some of the growth laws. By treating the main contents of a cell as discrete variables, the model has produced cell-to-cell variability in division times with distinct dormant states. The occurrence of long division times was shown to be perturbed by a transient downshift and the resource allocation strategy.

The model illustrated how a dormant subpopulation could comprise various physiological types, such as a collapse in translation or metabolism/energy production and how these might be interconnected. In this model, a collapse in energy production (i.e., hitting *m* = 0) was almost a necessity to reach a collapse in the translation system (*r* = 0). The existence of different classes of dormant states and their correlation is, in principle, an experimentally testable prediction, though it requires the development of single-cell experimental methods that can tell which part of the growth process is corrupted.

Consistent with existing literature [14, 16, 38, 40], we observed that the growth structure, i.e. lineage versus branch growth, strongly affects the occurrence of the dormant phenotypes. Growth in the branch structure led to much fewer cells in the dormant state. The effect was strong because, in the current model, there was a strong correlation for a slow-growing cell to produce another slow/dormant progeny.

Our model predicts the memory of dormancy, where the appearance of the dormant cell is preceded by longer inter-division time in the lineage (Fig. 5BCD.). It is worth noting that literature that studies bacterial ageing points to the existence of memory [47–49], though mechanism has been mostly attributed to the asymmetric division of damage aggregates that stresses a cell [50], and they may also compete against ribosomes for space [51]. Our current model assumed equal cell division as much as possible, and we still observed the dormancy because of the positive feedback. It is an interesting extension of the current model to include the effect of asymmetric division and damage aggregates.

Assuming that the dormant population is also a persister population, this model provides qualitative explanations for a few key results obtained in a previous experimental study on bacterial persistence [6] as follows.

First, the model reproduced persistence as a growth-rate-dependent phenomenon: There is a clear correlation that the higher the growth rate, the lower the dormancy fraction. The spontaneous persisters were below detection in the fast-growing cells in a rich medium[52], but in an exponentially growing population in a minimal media at an intermediate growth rate significant number of persisters were observed [6]. The simulations in this study suggest that spontaneous persister only exists at high enough frequency at slower growth rates, at least if a collapse in the central metabolism forms these persisters. According to our model, this observed growth rate dependence can be explained by the nutritional and energy state of the cell being closer to a collapse at lower growth rates. Here, it is worth mentioning that the growth rate is not the sole determinant of the dormant cell frequency; how close a cell is to a collapse depends also on which parameters control the growth rate.

Second, it is worth discussing the downshift in the current model and its relation to the experimental observation. In the current model, downshift induces a higher level of dormant cells for the intermediate to high growth rate condition, because the cells keep dividing even though the external nutrient is suddenly depleted. In the case of *E. coli*, wild-type cells would show stringent response in such a sudden downshift, where the alamone (p)ppGpp accumulates quickly [53, 54]. (p)ppGpp then triggers the regulatory mechanism to halt the growth to prepare the cell for starvation. Hence, the current model may be more appropriate for (p)ppGpp deficient, so-called relaxed mutants. In ref. [6], a mutant deficient in (p)ppGpp production (*relA*^*−*^ strain) was studied. It has been observed that the strain had a higher general level of persisters in the steady state growth, but upon the sudden downshift, it showed a factor of 10 to 100 more persisters for 2 to 7 hours after antibiotic application, with more persisters for a higher growth rate. This is qualitatively consistent with the current observation. It should be noted that, in the longer time scale (a day or more after the downshift), the downshift effect disappeared in these mutants [6], indicating the possibility that the long-term persisters are formed in a different mechanism.

The molecules (p)ppGpp are both involved in finetuning of physiology during balanced growth but are also necessary to elicit various stress responses during entry into a stationary phase, which is a qualitatively different state than the balanced growth state [53]. The model presented here does not consider these types of explicit bacterial stress responses. However, stress responses likely play a key role in physiology during a downshift for the wild-type strain. It is known that wild-type cells undergo reductive divisions during nutritional stress and that these lead to a reduction in size. In this process, all DNA replication will finish, and the cells will divide until one or two chromosomes per cell [53, 55]. Because faster-growing cells contain more origins of replication, which leads to more chromosomes when DNA replication is finished, they experience more reductive divisions and hence have a bigger probability of a key cellular component going below a certain threshold that requires a significant amount of time to regrow. We have shown in the Appendix how a simple model of reductive division can predict a strong effect of a transient nutritional downshift, as an alternative to the main model presented here. The main prediction from the reductive division model is that the persister fraction should increase with the downshift period until it saturates at a level, where all cells had time to go through the total number of possible reductive divisions.

Before concluding, we emphasize that our model is based on the well-characterized phenomenology of medium to fast-growing bacteria to study how dormancy could appear in a sub-population. However, we did not take into account the inactive ribosomes, ribosome degradation and turnover, which increase when nutrients are limited [56, 57], and the degree of increase depends on what is limiting (e.g. carbon source or phosphate source) [2, 58]. We did not consider inactive ribosomes and turnover since their roles and regulations are unclear, but as better understanding becomes available, it will be fruitful to extend the current model to take them into account. Including ribosome degradation kinetics upon starvation will be especially relevant to analyzing long-term starvation.

All in all, this study adds to a coarse-grained understanding of ffluctuations in bacterial physiology. Assuming that the machinery central to metabolism, translation, and cell division has a “threshold” below which the cell growth is strongly impaired, rare but distinct dormant states of a cell are expected. Despite the simplicity, this model provides qualitative tendencies consistent with the experimental observations, providing a baseline to discuss dormancy and persistence occurring without specific molecular mechanisms together with other works looking for a robust and universal explanation of the phe-nomenon [22, 37, 59].

## ACKNOWLEDGMENTS

MSS and NM were supported by VILLUM FONDEN (00028054). NM was supported by Novo Nordisk Foundation (NNF21OC0068775).

## Appendix The reductive division model

In the following, we present an alternative model, which includes a stress response. We want to demonstrate that such a model can arrive at qualitatively similar results as our main model, but it also leads to differences in the model predictions.

The model is based on reductive division. When *E. coli* is growing exponentially, the number of chromosomes is positively correlated with its growth rate, such that as the population grows faster, the average cell will contain more chromosomes. If they run out of some key nutritional component, they will enter a stationary phase, and with that comes a number of reductive divisions. The population does not increase its biomass, but the cell number still increases, meaning that cells are not growing, but still dividing and consequently reducing their size. Cells growing exponentially can have more than 6 chromosomes per cell [30], which means they will go through 2-3 divisions before they reach 1 chromosome. The assumption is that these cells do not produce new biomass so that their cellular content will be divided between the new smaller cells, which would increase the probability of some cells stochastically going below a threshold. The model assumes a key protein *x*, where the lack of it in a cell induces a persister state. This idea is illustrated in Fig. 7.

**FIG. 7.**
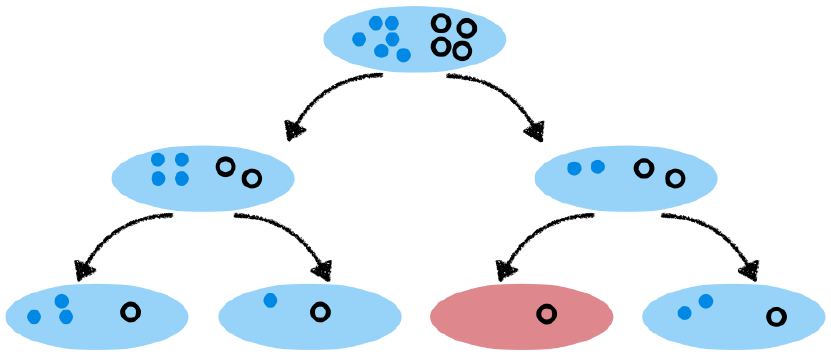
Reductive division during a stress response. Since the cells divide until they contain only one chromosome, they will also distribute other cellular content without potentially producing too much new protein.

At the starting point of the downshift, we assume that each cell contains *x* proteins. This could be a random variable, but let’s for simplicity assume that this is the same value for each cell. At each division, *x* is divided between the daughter cells following a binomial distribution, with *x*_1_ being the value in one cell and *x−x*_1_ in the other cell: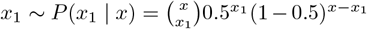. Now this process repeats a few times, depending on the initial number of chromosomes per cell, until the cells reach 1 chromosome. A binomial random variable conditional on another binomial random variable, is itself a binomial random variable, with the probability for one protein being in a specific cell *p* = 0.5^*σ*^, where sigma is the number of divisions. Thus, we arrive at the probability distribution for cellular content of *x*_*ds*_ after *σ* divisions:

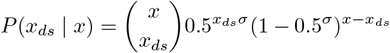

The number of divisions *σ* is approximated by the mean number of origins per cell, which is given by 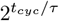, where *t*_*cyc*_ is the mean genome replication time and *τ* is the interdivision time. The cell then needs to divide 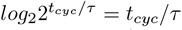 times to reach one origin per cell. So that *σ* = *t*_*cyc*_*/τ* .

The probability to reach *x*_*ds*_ = 0 is given by 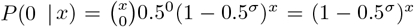. If we then compare the probability of reaching the zero-state with and without the downshift it depends exponentially on the number of reductive divisions and consequently on the initial number of chromosomes, which is given approximately by *σ ≈ log*_2_(*N*_*chromosomes*_). As an example, let’s compare the difference between 1 and 2 reductive divisions, corresponding to either 2 or 4 initial number of chromosomes:

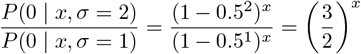

As *x* increases from 0, this number increases exponentially. The downshift thus has a much bigger effect on a population of faster-growing cells, compared to a population of slower-growing cells.

It is not easy to determine the dynamics of this process. There are several sources of noise, such as cell age, replication time, cell-to-cell variability of content, etc. However, we can expect the probability of entering a zero-state to increase rapidly in the initial part of the downshift and then saturate, as all cells have gone through all the reductive divisions.

Given that cells perform more reductive divisions if they contain a higher number of origins prior to the stringent response, faster-growing cells, with more origins, will see a bigger effect of a downshift than slower-growing cells, with fewer origins per cell. This qualitatively explains the higher fraction of dormant cells in the faster-growing population.

Next, we extend this consideration to analyze the time dependence of the dormant fraction of the cells upon downshift. Here we assume a very simple model where the cells go through reductive division following a Poisson process. We further assume that the fraction of cells to have undergone the first division increases with a rate *λ*, and then the next division happens with the same rate *λ*. This is illustrated for two divisions in the following, where the index corresponds to the number of divisions

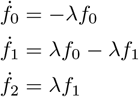

These equations have the following solution, with initial condition *f*_0_ = 1 and *f*_1_ = *f*_2_ = 0.

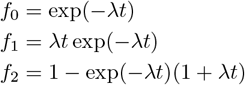

Since these three variables sum to 1 at all times, they correspond to fractions. Thus they are the probability of a cell having undergone 0,1, or 2 divisions following a downshift. These fractions can now be used to describe the probability of a cell reaching a zero-value state, assuming that the initial value is x. The probability to reach a zero state increases with the number of cell divisions, which means that cells of the subpopulation *f*_2_ have a higher probability of reaching zero than the *f*_1_ population. The *f*_0_ has no probability to reach the zero-value state since the cell experiences no dilution. In the preceding section, the probability of a zero state was shown to be *P* (0 | *n*) = (1 *−* 0.5^*σ*^)^*x*^. Thus the probability for a fraction of the population to reach zero can be described as the product of each population fraction with their corresponding probability of reaching the zero state:

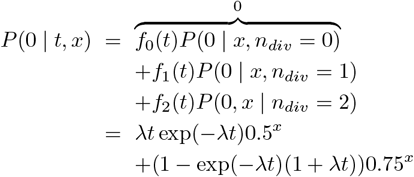

The time course of this probability is shown in Fig. 8 with *λ* = 0.1. We observe a rapid initial increase followed by saturation.

**FIG. 8.**
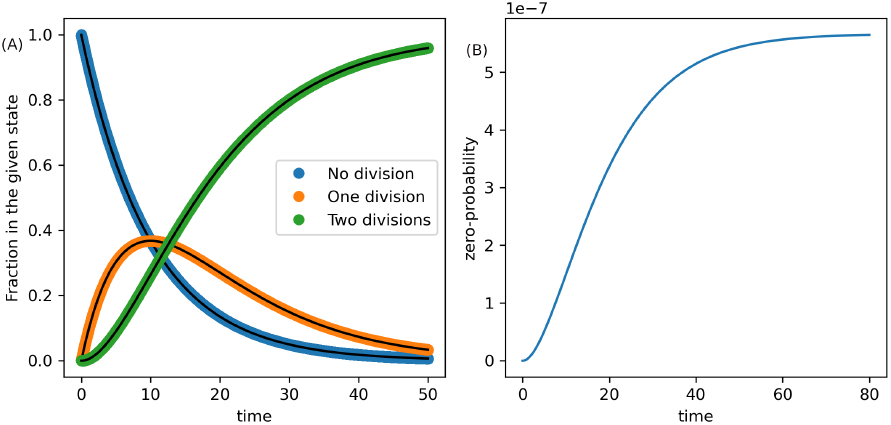
(A) Probability of either 0,1, or 2 divisions. (B) The probability of reaching a zero-state, which in this model is assumed to be dormancy. There is an initial increase in the probability, as the cells start to divide, but the probability saturates, as the cells enter the dormant state. *λ* = 0.1 and *x* = 50 were used in the plot.

