## Supplementary Material for "Simple bacterial growth model for the formation of spontaneous and triggered dormant subpopulations"

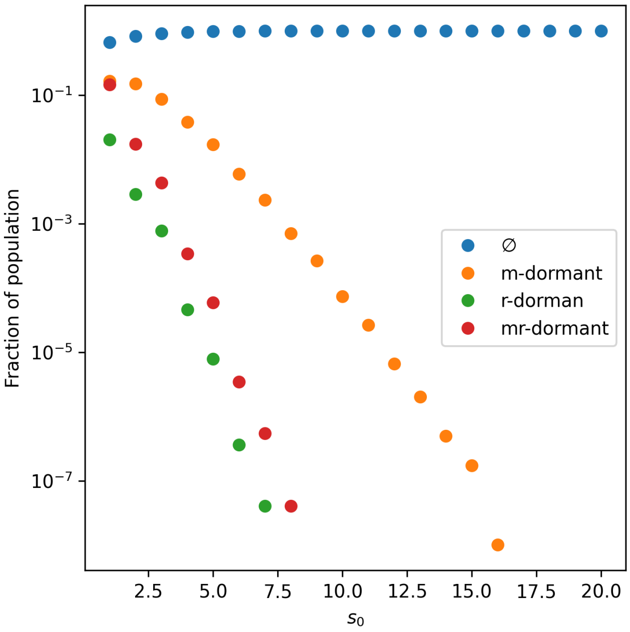


FIG. S1. The effect of varying the threshold for cell division parameter $s_0$. $\emptyset$ indicates the non-growring population. The higher the level of $s_0$, the lower the probability of reaching a dormant-state due to relative noise in cell division.

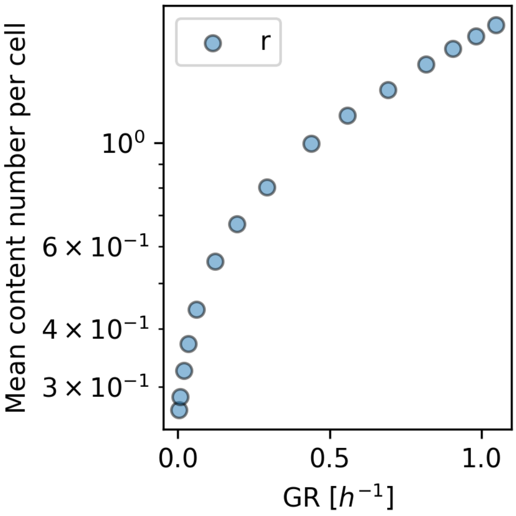


FIG. S2. Growth law depicting the ratio of $r$ in the total proteome as a function of growth rate. This ratio is given by $r/(s+m+r)$.

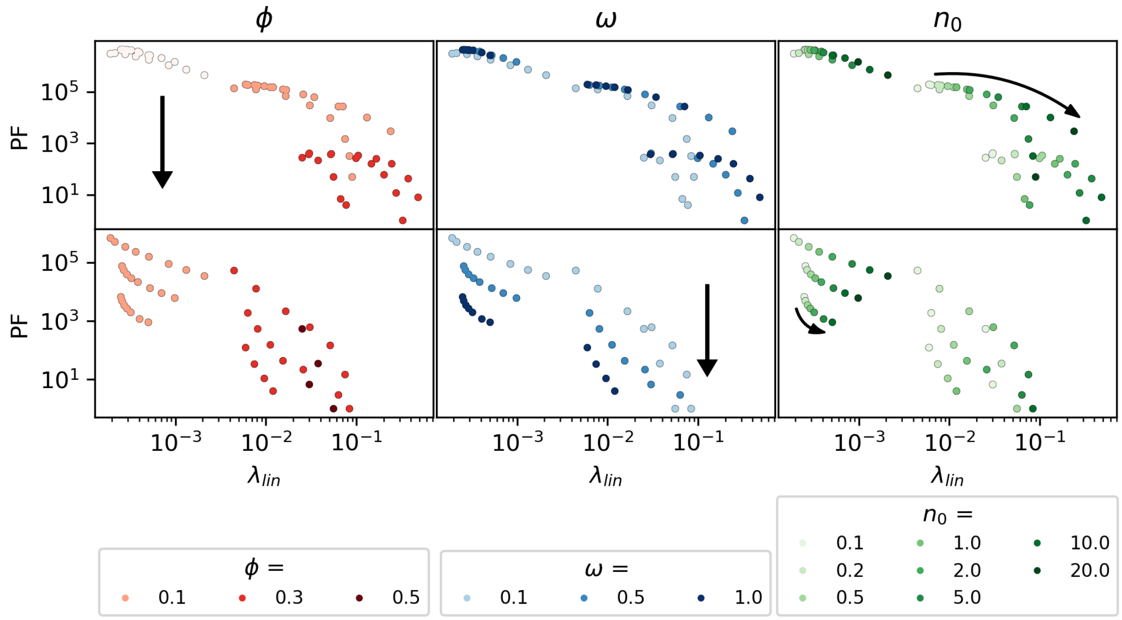


FIG. S3. Growth rate dependence of the dormant cell fraction. The holizontal axis shows the lineage averaged growth rate $\lambda_{lin}$, and the vertical axis shows the number of $m-$ dormant (top panel) and $mr-$dormant (bottom panel) cells per $10^9$ cells in lineage growth. The color indicate the parameters, to show that the dormancy is not just a function of the growth rate but it also depends on how the growth rate is varied.

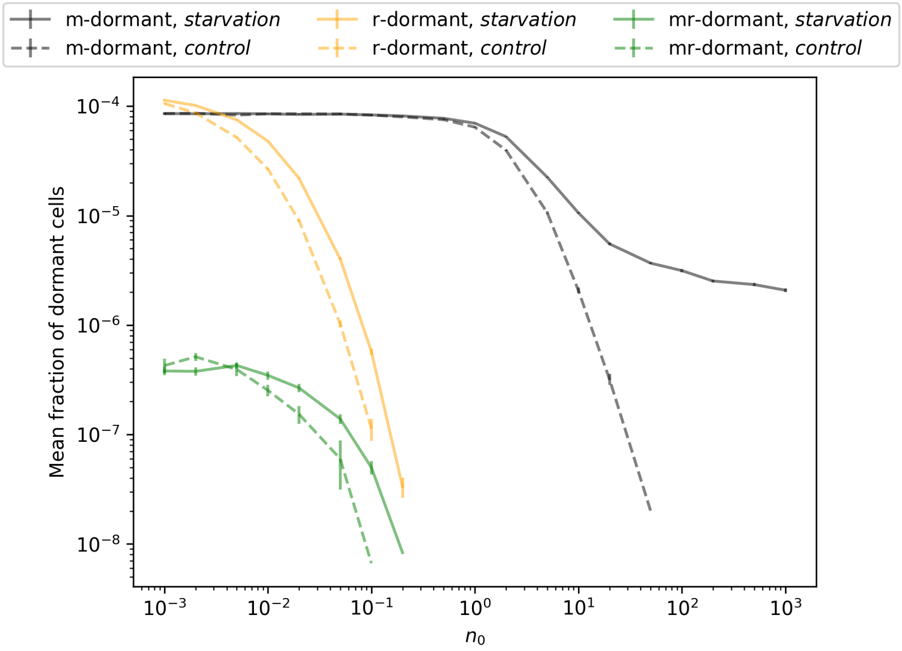


FIG. S4. The effect of feast-famine cycles on dormancy. The nutrient is kept to the value $n_0$ (the horizontal axis) for the control. When the feast-famine cycle is applied, the value of $n_0$ in the feast period is shown in the horizontal axis, which lasts 10-time units, and the starvation period of $n_0 = 0$ lasts 10-time units, and this feast-famine cycle is repeated. The vertical axis shows each dormant cell's mean fraction. The error bar shows the standard error of the mean from independent simulations.
